# Social anchors: how previous social locations shape current animal movement and habitat selection

**DOI:** 10.64898/2026.10.16.759611

**Authors:** Jillian M Kusch, Jack G Hendrix, Frankie D Tu, Kristy M Ferraro, Quinn MR Webber, Eric Vander Wal

## Abstract

Memory of past space use is an important factor in habitat selection, but memory of what? The concept of memory captures the geographic spaces that an animal has used in the past. There are, however, competing reasons why animals occupy a given space. Species may occupy space due to particular aspects of their memory. Using integrated step selection analysis, we tested the relative importance of memory: previous use of high forage quality or where social interactions occurred at two temporal scales in caribou (*Ranger tarandus*). The most important parameter to explain intra-seasonal movement was geographic memory. At the interannual scale, caribou more strongly revisited locations with high quality forage and where they had formed dyads with conspecifics. We posit that remembered locations of social fusion can function as social anchors. The effects of memory on movement behaviour depend upon the context—both social and resource-based—and not geographic space alone.

## Introduction

Habitat selection describes the choices that animals make among available environments (1,2). The decisions that govern animal movement are dynamic and can vary across individual preference, seasons, life stages, and population dynamics. The factors that are most often deemed to affect habitat selection are external environmental variables (i.e., resources, risks, and conditions; (3)) such as food availability (4), shelter (5), thermal conditions (6), predation risk (7), and reproductive opportunities (8). However, more recent research has suggested that previous experience and familiarity, often described as memory, is a critical component of movement and space-use decisions (9,10). Spatial memory enables animals to store and retrieve information about the locations of resources, risks, and conditions, allowing them to make informed movement decisions based on past experience rather than solely on immediate environmental cues (11). Ravens (*Corvus corax*) have demonstrated using memory to return to previous wolf (*Canis lupus)* kill-sites in contrast to following cues of current wolf presence (12). Further, memory of migration routes is more effective than perception of current resources in directing individuals to seasonal ranges (13). These examples highlight memory as a fundamental driver of habitat selection and movement, particularly in heterogeneous and temporally variable landscapes.

Rather than assuming individuals have complete or instantaneous knowledge of the landscape, memory-based habitat selection models recognize that animals gradually acquire spatial knowledge through exploration and repeated use of an area, forming an internal representation, or *cognitive map*, of their environment. An organism’s cognitive map can range in temporal scale from long-term memory with seasonal or yearly patterns to short-term memory focused on locations visited recently. Including both short-term diurnal memory and long-term seasonal memory contributed to improved habitat selection model performance for feral hogs (*Sus scrofa;* (14)).

Although memory has become an increasingly important component of habitat selection and movement models, it is often represented in a highly simplified manner. In many memory-based frameworks, an animal’s previous locations are incorporated as a spatial memory surface, where revisitation probability is assumed to reflect familiarity with a location. These cognitive maps can be represented as home ranges (2), residence times (15), time since last visit (16), and occurrence distributions (14), but all are configurations of the geographic space—a series of XY coordinates. While these approaches successfully capture broad patterns of site fidelity and recursive movement, they generally treat all visited locations as equally informative, regardless of the ecological context in which they were experienced. As a result, the memory term often reflects only where an individual has been, rather than what it encountered there. Geographic space and environmental space are intrinsically linked (sensu van Moorter (2)), and it is likely that by representing memory as only a spatial map, that the environmental context is missing. Locations associated with abundant food, successful reproduction, predator encounters, human disturbance, or poor foraging conditions would contribute similarly to the memory surface despite having very different behavioral consequences and requiring different interpretation. Studying memory as geographic points overlooks the possibility that animals develop memories linked to the quality, predictability, and outcomes of previous experiences, rather than simply retaining a record of geographic coordinates. Incorporating contextual information into memory-based habitat selection models has the potential to better represent how animals use experience to guide future decisions, offering a more biologically realistic framework for integrating past experience into movement and space use in heterogeneous and dynamic environments.

Species’ traits also influence both habitat selection and the context of their memory-based spatial maps (17). Specifically, social species are known to use the presence of conspecifics to inform movement (18). Increased foraging success (19), exploration of risky places (20), and predator avoidance strategies (21) have been attributed to socially-driven movement behaviour. Similarly, social species may be more likely to use social memory that influences movement. We define social memory as an individual’s ability to recognize and remember other individuals, social relationships, or locations associated with previous social interactions, enabling movement decisions to be shaped by the social landscape. Current conspecific proximity (22), group size (23), and local density (24) impact individual habitat selection, and therefore it is likely true that an updated reference of social memory is also crucial in habitat selection. Integrating social memory into habitat selection analyses offers an opportunity to better understand how animals navigate both their physical and social environments and how these processes jointly shape movement and space use. Memory of the social and environmental landscapes work collectively in habitat selection, suggesting that considering both is required for a better understanding of habitat selection. Bison (*Bison bison*) with increased social familiarity rely on their conspecific movements to inform their own movement, while those with increased environmental familiarity use their own previous movements to make future decisions (25).

Woodland caribou (*Rangifer tarandus;* hereafter ‘caribou’) make movement decisions based on both the social and environmental landscape (26–28), and therefore are an excellent model species for disentangling the variables that contribute to memory. Caribou are a fission-fusion species that forms small groups with variable membership over time. During the winter they aggregate into larger herds, likely for increased foraging success (29). Nutritional access is also increasingly limited in the winter and caribou are also known to select habitats with increased lichen % N in winter (30). Caribou also make movement decisions that are informed by long-term environmental memory and sensory inputs from their surroundings (31). Therefore, expectations for how geographic, social, and environmental spaces influence movement can be examined.

Our objective is to test how components of memory, namely social and nutritional memory, affect habitat selection in comparison to the general geographic memory paradigm. We also investigated how these forms of memory influence habitat selection at both a long-term and short-term scale. We hypothesized that social and nutritional components of memory would have a stronger effect on movement than geographic memory at the long-term scale. We predicted that caribou would select for areas of previous social encounters and areas of higher nutritional quality, within their winter seasonal range. We hypothesized that caribou would avoid areas associated with short-term memory and that geographic memory would be the strongest predictor for movement at the short-term scale due to depletion of resources on the patchy winter landscape. At the short-term scale, avoidance of depleted resources may surpass their selection for social interactions within these areas. Our research aims to disentangle the geographic, nutritional, and social aspects of memory and contribute to both the memory-movement and social-spatial interfaces.

## Methods

### Study area

We conducted our study on Fogo Island, a small (∼238 km^2^) island off the northeast coast of Ktaqmkuk (Newfoundland [NL], Canada). The habitat types on Fogo Island include lichen barren-grounds, scrubby boreal forests, and boggy wetlands. The NL provincial government introduced woodland caribou to the island in the 1960s (32). The population reached a peak of about 300 individuals in 2018-2020 (33) and have since declined to about 150 individuals (30). This study system is a long-term ecological project that began in 2016 and continues to monitor adult and calf survival, movement patterns, and the nutritional landscape that underpins it all.

### Caribou GPS data and iSSA preparation

Between 2016 – 2023, we collaborated with the NL Wildlife Division to deploy GPS collars (Lotek Wireless Inc., Newmarket, ON, Canada) on adult female caribou (2016-2019 deployments GPS4400M, 1250 g; 2022-2023 deployments Litetrack Iridium 420, 440g). Capture and collaring procedures were conducted by Wildlife Division staff following government protocols. We prioritized re-collaring the same individuals over time where possible to collect multiple years of data on the same individual. We set the deployed GPS collars to record XY coordinates every 2 hours. A mean of 24 individuals had collars per year (minimum = 14; maximum = 30). Not all collars resulted in usable data due to lost signals or improper settings resulting in between 1 and 22 individuals with usable data per year.

We used integrated step selection analyses (iSSAs, (34)) to test our memory-based hypotheses. We calculated the Euclidean distance in meters (termed ‘step length’) between consecutive GPS points to generate ‘steps’, and the change in direction (termed ‘turning angle’) between steps. We modeled turning angle with a von Mises distribution where 0 represents straight-forward movement, and 180° represents a complete reversal of direction. We then created random steps for which to compare the “true” steps against for testing selection or avoidance of memory. We used both step length and turning angle to create biologically realistic distributions from which to draw the random steps. We generated n = 10 random steps for each “true” step. In a sensitivity analysis, 10 steps are deemed to be sufficient to convey the variation in generated random steps (35). For each true step, and the corresponding 10 random steps, we created a shared numeric ID to identify each set of steps for modeling purposes.

Using the coordinates from the GPS data and the calculated step lengths, we cleaned the data before modeling. We removed any GPS data that were likely erroneous, e.g., coordinates in the ocean or step lengths that exceeded 30 km due to being biologically impossible for Fogo Island’s size of 18 km in length. After cleaning, we had 408,788 steps from 47 individuals. Caribou tend to be more social in winter than summer in our system (33). We excluded summer from our analyses, as very few social encounters occurred (n = 3/individual/summer/year), and a social memory layer could not be created with enough variance to converge (Suppl Mat. Figs S1,S2). We defined the winter season as 1 January – 1 March. This period represents where there is continuous snow cover > 75 % across years (30,36). For the winter period, we had n = 62,022 steps across n = 43 individuals (and 93 individual-years). We further reduced the dataset for the long-term memory model that required sampling from two winter seasons in a row. 32 individuals had repeated samples that could be used for inter-seasonal modeling with n = 46,156 steps. We then constructed layers on which to model the movement data. We created geographic, social, and nutritional memory layers at both the long-term and short-term scale.

### Long-term memory layer

We built long-term geographic memory layers using the GPS fixes from the previous winter for each caribou. We generated utilization distributions (UD) with the kernelUD function of the adehabitatHR package (37). Both utilization and occurrence distributions have been used in habitat selection modeling (38), but we chose to use UDs over occurrence distributions to avoid performing interpolation on which additional models were conducted. Further, animals that portray significant site fidelity to an area may have occurrence distributions that are vanishingly small and not capture their true distribution (39). For each UD, we generated an isopleth to determine values of recent, repeated use. Values of 0 indicate coordinates that were heavily visited in the previous winter and therefore represent increased memory, while values near 1 indicate coordinates that the caribou have not visited and have no memory of from the previous winter. For each unique step and the corresponding randomly generated steps, we extracted the value of the isopleth at each step’s coordinate. We also built long-term geographic memory models for UDs restricted with a kernel of 50 % and 25 % to check for robustness to our method. We found similar trends for these reduced UD sizes and have presented those results in the Supplementary Materials (Figs S3,S4).

### Short-term memory layer

Recent memory is commonly represented as a dynamic spatial layer that changes over time according to an individual’s movement history (10). The changes over time can be modeled as a decay function (40) or through restricting the temporal period to represent memory (14). We opted for restricting the temporal period to represent memory, where we built short-term geographic memory layers for each step taken for each caribou in the winter period. We used the ‘kernelkcbase’ function of the adehabitatHR package (Calenge 2006) to create UDs. The function allows for smoothing in both temporal and spatial parameters. We smoothed the UD with a 14-day temporal window and a 500 m spatial distance. We chose 500 m as caribou are shown to have a perceptive limit at that distance (31). For each UD built, we applied the same isopleth method as from the long-term memory step. For each unique step, we extracted the value of the isopleth at each step’s coordinate, and at the corresponding randomly generated steps. We also completed an analysis with a 3-day temporal window to determine how sensitive our temporal window choice was to changes in overall trends. We did not observe a difference in trends between 3 and 14 days. Therefore, we’ve chosen to present the 14-day models in the main text and the 3-day results can be found in the Supplementary Materials (Fig S5).

### Social memory layers

We used previous ‘fusion’ events to represent social memory. Fusion indicates places where individuals are likely to want to form groups with conspecifics for some social benefit. We denoted fusion events based on previous caribou fusion literature (26). Briefly, we determined fusion locations based on coordinates where caribou were within 50 m within 5 min of each other, using the ‘spatsoc’ package (41,42). Caribou had to remain within 50 m of each other for 2 subsequent time-steps to be considered a fusion event. A total of 11,720 fusion events occurred during the winter season across the study period. For each individual, we generated maps of coordinates for fusion events restricted temporally for both the long-term and short-term models. We calculated ‘distance-to’ the closest fusion event within the restricted period for each step taken by a caribou, and the corresponding random steps (43).

### Nutritional memory layers

We used a stoichiometric distribution model (StDM) of % nitrogen (N) for *Cladonia* lichen, derived from Hendrix *et al*. (30). Briefly, we collected vegetation samples of *Cladonia* lichen, which we then dried, and weighed, and calculated %N for each sample. We regressed topographic predictors across Fogo Island with the %N values for each sample to build a StDM. Our StDM predicted surfaces were calculated at the finest scale of our set of predictors, which was 5m x 5m from the digital elevation model. We aggregated %N values to a 30m pixel, to match the land cover cells used to predict habitat selection in this population (27). Some locations, such as those within bodies of water and on roads, had missing values from the StDM as we did not predict %N into these land cover types. When summarizing %N, we treated these missing values as zeros to reflect the fact that they do not offer any forage opportunities. We used the map as a fixed value for nutrition as relative differences in %N in space have been shown to not change drastically across years (44). Further, Thompson *et al*. (16) reported increased model performance with fixed resource layers as animals may remember these resource layers even outside of their ‘seasonal’ window.

We used the UDs at the short-term and long-term scales with the “mask” function of the raster package to create nutritionally-informed memory maps (45). We assigned all values outside of the UD to zero to represent no memory of the nutritional landscape beyond the memory region. Using these nutritional memory maps, we extracted the *Cladonia* %N value for each step taken by a caribou, and for the corresponding random steps.

### Land cover layer

Recent work has demonstrated that including elements of the physical environment are critical when conducting analysis on social data to avoid spurious results (46). We included forest cover in our model as previous work has shown that caribou select for open habitat during the winter season (20). For each 100×100 m cell we calculated the proportion of forest cover based on a dichotomized land cover class of ‘forest’ and ‘open’ from the Canadian Forest Service system (47). Because we know that this population of caribou avoids forest in winter, the inclusion of forest proportion in the model also served as a ‘check’ for model performance within the analyses.

### Data analysis

We constructed two iSSA models, one to test long-term memory and one to test short-term memory. Both models included proportion forest cover, geographic memory, social memory, and nutritional memory. We also included random effects of year and the numeric ID that connected the ‘true’ step and its random steps, along with a random intercept and random slopes for individual ID following Muff *et al*. (48). We fit our models as a mixed Poisson regression, with a binomial 0/1 response of randomly generated vs. true steps. We completed correlation tests between the nutritional landscape and locations of fusion events to assess collinearity.

To interpret the results of our iSSA models, we calculated relative selection strengths (RSS) for each parameter of the model (34). An RSS is a representation of selection and avoidance for a given parameter when all other parameters are held at their mean. The logRSS can be plotted as a function, where values > 0 indicate selection and values < 0 indicate avoidance, relative to a reference value for the parameter of interest. We used the mean/median value of each covariate as the reference level. The slope of the RSS indicates the strength of the selection and avoidance. We also calculated RSSs by year for each model, and have included those in the Supplementary Materials (Figs S6–S13).

We evaluated goodness of fit of our iSSA models using a comparison to null models with a log-likelihood ratio test. Our null model contained only the proportion forest parameter, and the interaction with Animal ID. We removed all parameters that involved memory as these were the parameters of interest. All analyses were conducted in R v4.3.3 (49) in a targets workflow (50), using packages including amt (51), data.table (52), ggplot2 (53), glmmTMB (54), sf (55), and terra (56).

## Results

Both the short-term and long-term models performed better than our null model that included only proportion forest and an interaction with Animal ID. For the long-term model, we observed a significant improvement in model fit when adding memory parameters compared to the null model (−2ΔLL = 9243, df = 7, p < 0.001). For the short-term model, we also observed a significant improvement in model fit when memory parameters were added compared to the null model (−2ΔLL = 44580, df = 7, p < 0.001).

Overall, we found support for our memory-based habitat selection predictions in caribou (Table 1; Fig 1). For long-term memory, we predicted that caribou would avoid forested areas and would select for geographic memory. We also predicted that caribou would select for places where they had formed social groups in the previous winter (social memory) and patches with the highest nutritional quality (nutritional memory). As predicted, caribou avoided forest, with substantial individual variation, including one individual that selected for forest (Fig 1C). The RSS plot shows that caribou selected areas of increased memory in the previous winter (Fig 1A). However, the plot of used and available steps reveals that caribou spent most of their time in areas of decreased memory (Fig 1B). The selection for increased memory appears to be driven by the decreased selection of low memory compared to random steps, despite the overall use of low memory areas. Caribou also selected for areas where they had previously formed fusions with other conspecifics and areas of higher *Cladonia* %N (Figs 1D;1E). The correlation test revealed a slope of -0.16 between social and nutritional memory at the long-term scale, indicating that these forms of memory are distinct drivers of movement.

**Table 1.** Parameter estimates from long-term and short-term memory integrated step-selection analyses of caribou on Fogo Island, NL. The long-term model includes n = 46,156 steps from 32 individuals. The short-term model includes n = 62,022 steps from 43 individuals. Model summaries have been included for transparency, but for the purpose of interpretation, relative selection strengths (RSS) serve as a more relevant tool better (34).

| | Estimate ± SE | z | p |
| --- | --- | --- | --- |
| Long Term |  |  |  |
| Proportion forest cover (%) | -1.115 ± 0.115 | -9.66 | < 0.001 |
| 2 month inter-seasonal KDE | -0.790 ± 0.092 | 8.60 | < 0.001 |
| log(distance to fusion) | -0.219 ± 0.020 | -11.14 | < 0.001 |
| *Cladonia* % nitrogen | 1.211 ± 0.076 | 15.90 | < 0.001 |
| Short Term |  |  |  |
| Proportion forest cover (%) | -0.782 ± 0.057 | -13.63 | < 0.001 |
| 2 week intra-seasonal KDE | -3.374 ± 0.047 | -70.59 | < 0.001 |
| log(distance to fusion) | -0.299 ± 0.022 | -13.69 | < 0.001 |
| *Cladonia* % nitrogen | 0.726 ± 0.040 | 18.12 | < 0.001 |

**Figure 1.**
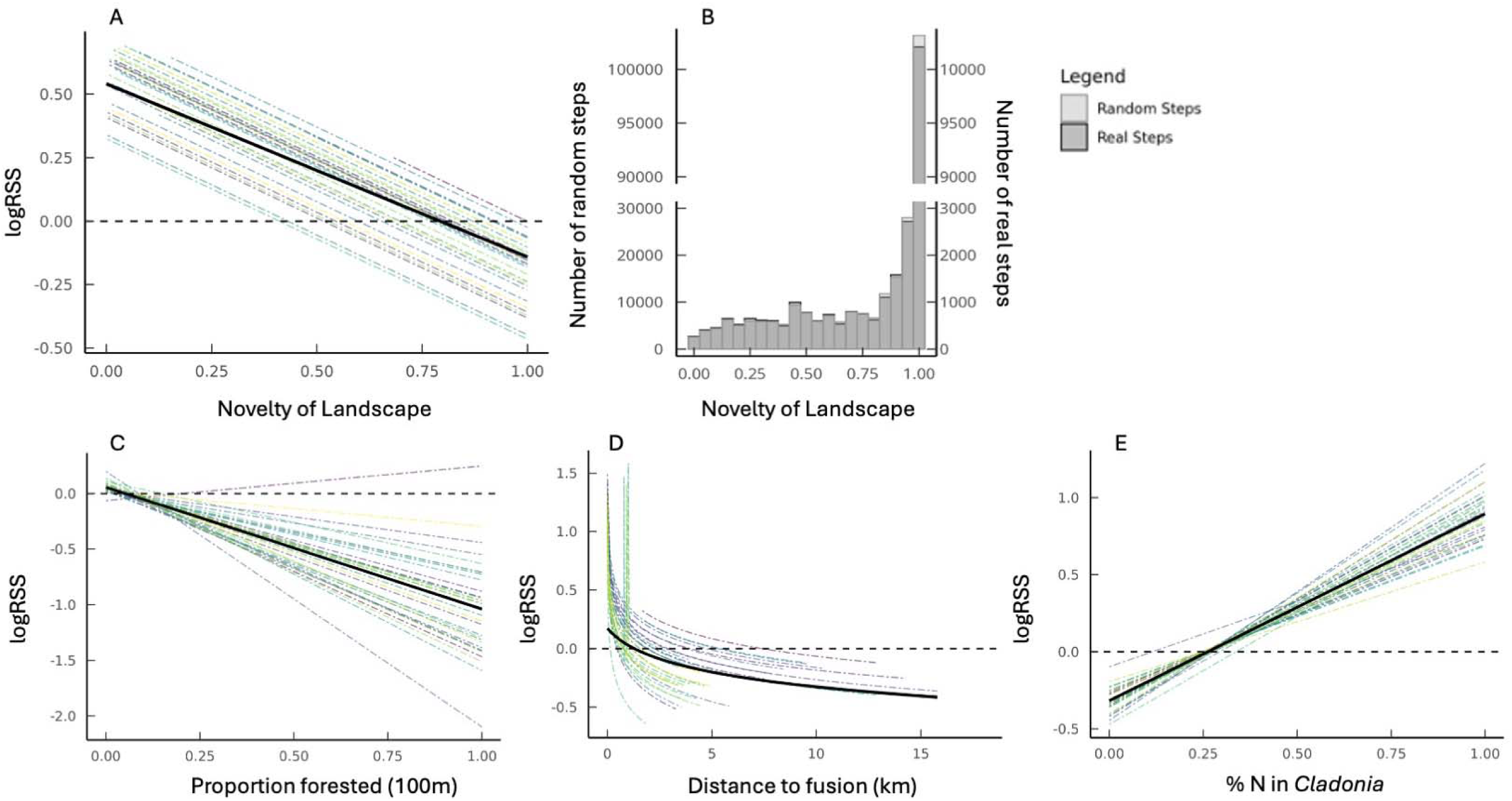
Model outputs for long-term memory iSSAs. Relative selection strength (RSS) plots for geographic memory (A), proportion of forest (C), distance to fusion (D), and % nitrogen in *Cladonia* (E). The thick black line represents the population mean selection strength. Each individual is represented by their own dashed line, showing variation within the population. A value above 0 on the y-axis (logRSS) indicates selection for each parameter, when all other parameters are held at the mean. (B) A histogram of real and random steps taken by caribou across all years, and their relative position within their geographic memory for long-term memory. Most of their time is spent in places within their range, but lesser used than the previous year. However, selection is for more familiar areas compared to random steps, as seen in A.

Our results did not support all predictions at the short-term scale (Table 1; Fig 2). We predicted that caribou would avoid forest at the short-term scale. Our prediction held, with all individuals selecting against forest (Fig 2C). We also predicted that caribou would avoid geographic, social, and nutritional memory at the short-term window due to avoiding areas of depleted resources. In contrast, we found that geographic memory had the largest effect on habitat selection, but caribou selected for areas from their most recent, familiar memory (Figs 2A;2B). Caribou selected for both past fusion locations (seasonal memory; Fig 2D) and *Cladonia* %N (nutritional memory, Fig 2E), but their effect sizes were much smaller than for geographic memory (Table 2). The slope for the correlation test between short-term social memory and nutritional memory was -0.09, suggesting that these parameters are distinct from each other in driving selection.

**Figure 2.**
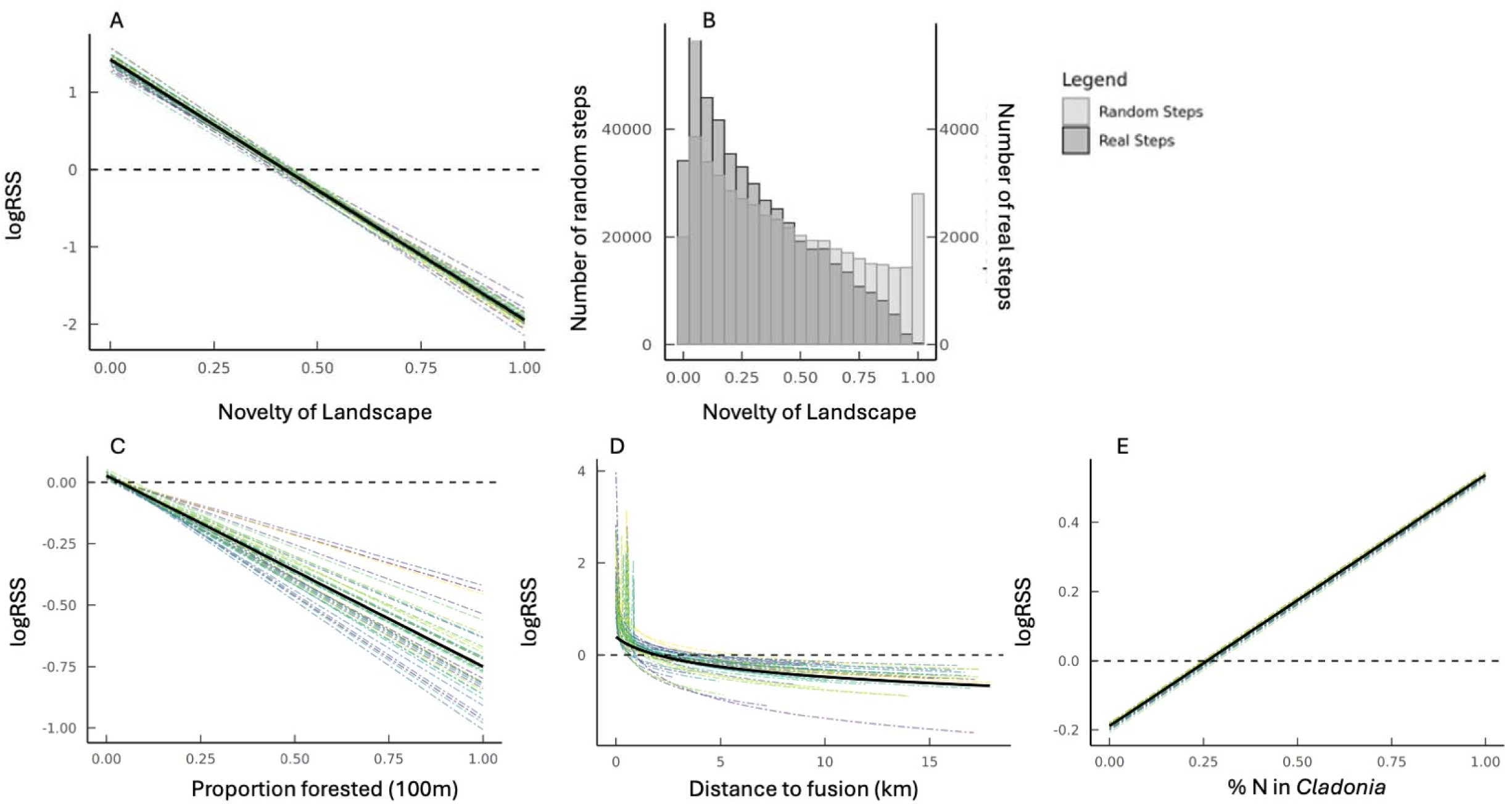
Model outputs for short-term memory iSSAs. Relative selection strength (RSS) plots for geographic memory (A), proportion of forest (C), distance to fusion (D), and % nitrogen in *Cladonia* (E). The thick black line represents the population mean selection strength. Each individual is represented by their own dashed line, showing variation within the population. A value above 0 on the y-axis (logRSS) indicates selection for each parameter, when all other parameters are held at the mean. (B) A histogram of real and random steps taken by caribou across all years, and their relative position within their geographic memory. Most steps occurred inside their short-term memory area, and selection for familiar areas was much higher for real steps compared to random.

## Discussion

We sought to evaluate memory within habitat selection models by comparing three contexts under which memory could be interpreted—geographic, social, and nutritional (Fig 3). Over the long-term, caribou used their seasonal memory to dictate habitat selection, but chose underused locations compared to the previous year’s locations. Importantly, social memory and nutritional memory were distinct drivers of habitat selection at this scale; caribou were not merely returning to previous geographic locations but responding to the contexts that were present in those locations in previous winters. At the short-term scale, caribou selected most strongly for areas they had recently visited (geographic memory) and responded less intensely to social and nutritional memory. Within recently visited areas, social and nutritional memory remained important, but their relative selection was diminished in comparison to just geographic memory. Memory has been studied across multiple temporal scales in animal movement, reflecting the capacity of animals to use both recent experiences and information accumulated over seasons or years to guide movement decisions. Our results suggest that for areas recently visited, the context of familiarity in that area is not yet learned. In contrast, individuals may develop familiarity with landscapes over longer periods with the context for those areas also being remembered. Feral hogs display strong selection for recent geographic memory, which is hypothesized to be due to uncertain resource availability where reliance on an environmental layer is likely less informative as the short-time scale (14). Selection for short-term geographic memory has also previously been shown in mule deer (*Odocoileus hemionus*), with recent experiences outperforming environmental factors in habitat selection models (40). Similarly, mule deer recent memory was also shown to decay quickly over time, suggesting that long-term memory and environmental cues become important as repeated use of an area occurs (40). Collectively, this work suggests that memory may operate simultaneously across multiple temporal scales. The persistence and decay of information determine how the context of past experiences and their relative importance shape movement decisions. The effect of memory depends not only on *where* an animal has been, but also on *when* it was there.

**Figure 3.**
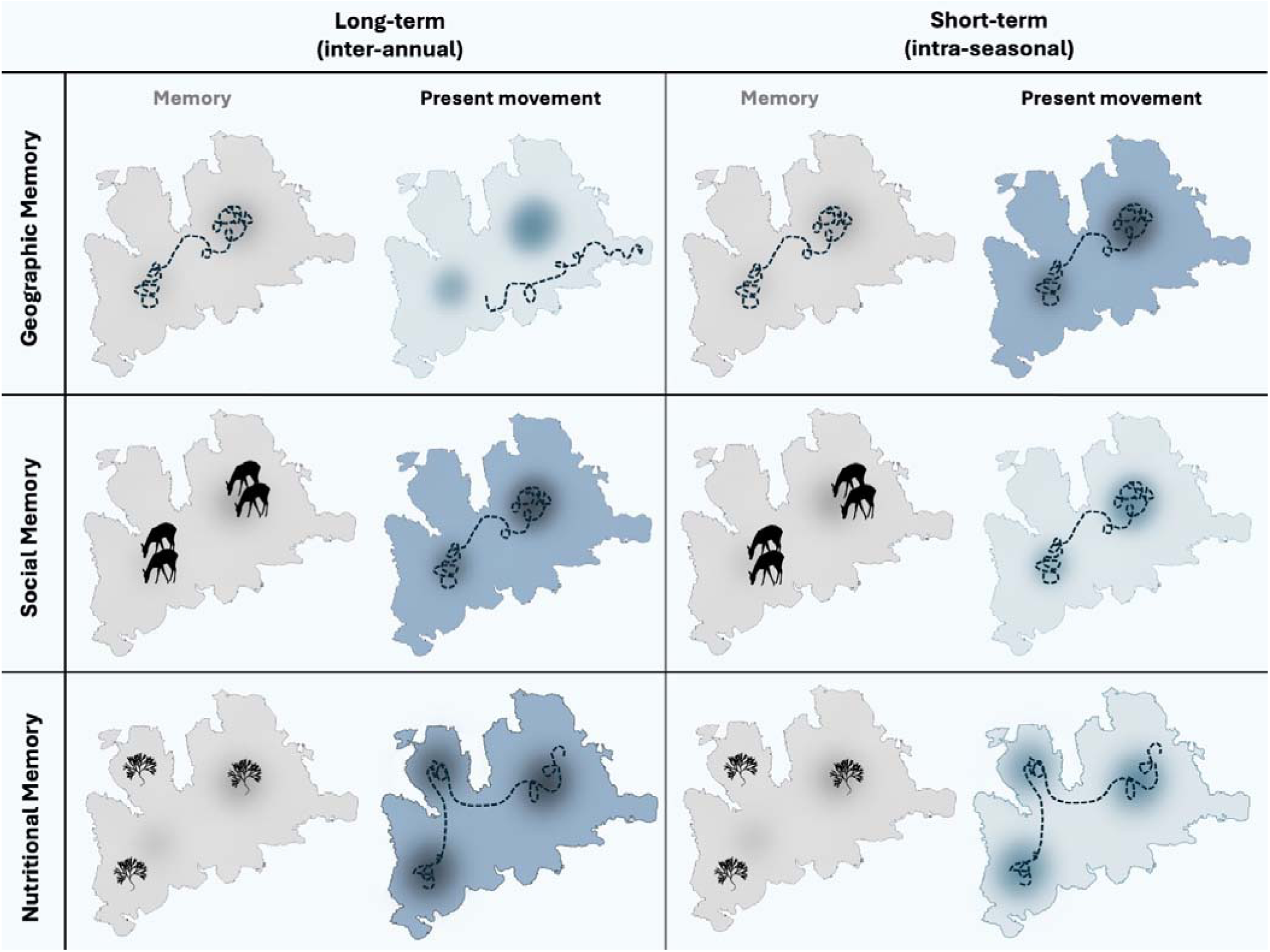
A conceptual summary figure of the results from caribou memory habitat selection on Fogo Island, NL. The dark spots on each map show the memory from the previous sampling period and caribou tracks either visit these spots or avoid them. Dark blue colours indicate which temporal scale each type of memory is most impactful. Silhouettes drawn by K. Ferraro.

The social context of memory was an important predictor of habitat selection for caribou as caribou selected for locations where they previously formed groups at both the short- and long-term scale. Selection for locations of previous grouping was particularly evident at the long-term scale. The long-term results suggest that caribou retain the memory of group formations upwards of a year after a given fusion event. We borrow the concept of a ‘spatial anchor’, which is a fixed landscape feature that anchors movement of an animal, to explain selection for previous social fusion locations (57). Spatial anchors guide movement as these locations contain vital features like dens, nests, or water sources. We propose that these previous social fusion locations serve as ‘social anchors.’ Rather than treating spatial anchors solely as features of the physical environment, socially-mediated anchors may emerge when individuals repeatedly associate particular locations with opportunities for social interaction or association. Social anchors are not mutually exclusive from spatial anchors, as some geographic locations have increased social interactions, like dens or water holes, but instead serve to describe locations where the social component might outweigh the environmental component. Through the process of returning to locations associated with sociality, geographic locations acquire social significance independent of their intrinsic environmental properties, becoming points around which movement and social behaviour are organized. Thus, social fusion may emerge from the interaction between a tendency to seek social partners and the capacity to remember where those encounters have occurred, creating feedback in which socially rewarding locations become increasingly salient and are revisited more frequently. Social fusion may therefore reflect not only the immediate convergence of individuals in space, but also a temporally extended process in which previous social encounters shape subsequent spatial decisions. Caribou may find that social fusions allow for selection of habitat that may otherwise be unavailable to them, like riskier, open habitat (26,35); the habitat that contains their winter forage—lichen.

Nutritional memory was an important parameter in habitat selection for caribou during the winter season. Other studies have demonstrated that animals return to previously visited resource-rich locations like forage patches or kill sites (58,59), but the inclusion of % N as a memory covariate appears to be unique to our study. Our results demonstrate that animals may not only remember resource-rich landscapes; they may also develop spatial memories of the specific nutritional characteristics associated with particular locations. As our study was completed in the winter, we also suggest that this selection for nutrient-based memory may be driven by the scarcity of resources on the landscape. When resources are patchily distributed, individuals that remember where profitable patches occur can reduce the costs of searching and preferentially revisit areas that have previously provided high energetic or nutritional returns (60). Caribou experience substantial spatial and seasonal variation in forage availability and nutritional quality, including variation in the availability of energy and protein among plant communities (61,62). Individuals may therefore develop an internal representation of the landscape that incorporates not only the location of forage patches but also their expected energetic or nutrient content even within a specific forage type like *Cladonia*. Further, cratering behaviour during the winter is extremely costly (63). Avoiding cratering in areas with poor forage quality might be a primary driver of remembering the nutritional landscape. Studying memory at such a nutrient scale of resource availability could be a novel step in determining how animals navigate heterogeneous landscapes and make repeated foraging decisions over time.

The concepts of site fidelity and memory are intertwined as they are both metrics of where an animal has been in the past, and their propensity to return to it (64). Across ungulates, species that display site fidelity are also more likely to rely on their memory in movement decisions (17). Previous literature supports that caribou, particularly woodland caribou, show fidelity to home ranges and, in some populations, return to the same areas for calving across successive years, although the strength of fidelity varies individually and seasonally (28,65). Our results agree with the general trend of site fidelity in that caribou selected for areas of increased geographic memory at both the short- and long-term scale. However, our results require some additional interpretation. At the long-term scale, while caribou remained in their seasonal memory throughout the sampling period, most steps occurred in the least familiar places, despite selecting for highly familiar areas (Fig 1B). We suggest that Fogo Island caribou do not prefer unfamiliar areas, but rather, they select habitat in a temporal cycle akin to nomadism (66). Indeed, in the winter season, plasticity around site fidelity has been observed as a flexible response shaped by seasonal climate conditions and individual experience (67). This perspective is particularly useful for understanding movement in heterogeneous environments where resource renewal occurs over long timescales, because it suggests that animals can use memory to regulate the timing of revisits.

## Conclusion

By incorporating different forms of memory into habitat selection analyses, our models provide a more mechanistic understanding of site fidelity (17), home range formation (68–70), migration (71,72), and recursive movement patterns that cannot be explained by environmental variables alone (60,73). While the inclusion of geographic memory into habitat selection models has shown improvement in model predictions (10,40), our analysis demonstrates that consideration for the context of that memory is also critical. The inclusion of social memory is especially essential for species where socially informed movement decisions contribute to site fidelity, collective movement, and the emergence of stable social and spatial structures. We have also extended the concept of a ‘spatial anchor’ to include places not just as features of the physical environment, but also as locations where the propensity for repeated social interactions occurs. An animal’s movement history is not simply a response to the current landscape but is also a response to its accumulated history of interactions with that landscape and the individuals within it.

## Acknowledgements

This work – data collection, analysis, and writing – occurred on the ancestral homelands of the Mi’kmaq and Beothuk. We thank Dara Albrecht for assistance in collecting plant samples, and the Government of Newfoundland for capturing and collaring caribou. A special thank you to the iSSA working group at Memorial University for troubleshooting and providing feedback to the many iterations of this project.

