## Supplementary Materials for "Social anchors: how previous social locations shape current animal movement and habitat selection"

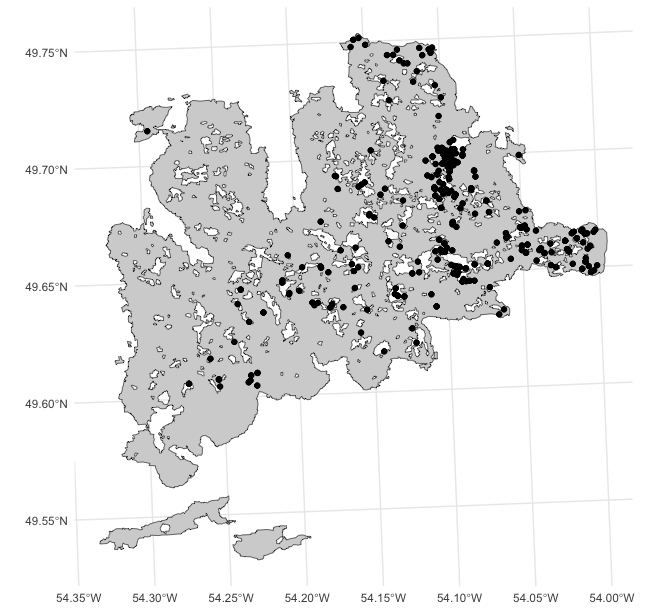

**Figure S1**. A plot of all fusion events for the summer period on Fogo Island from 2016 to 2025. Across all ten years (2016-2025), 334 fusion events occurred in the summer season (June 16 to August 16) involving 43 individual caribou. A mean of 15.53 fusion events for each caribou with a range of 1 to 62. A mean of 33.4 fusion events per summer, with approximately 1.5 events occurring per individual (as two individuals are involved in one fusion event) for each summer.

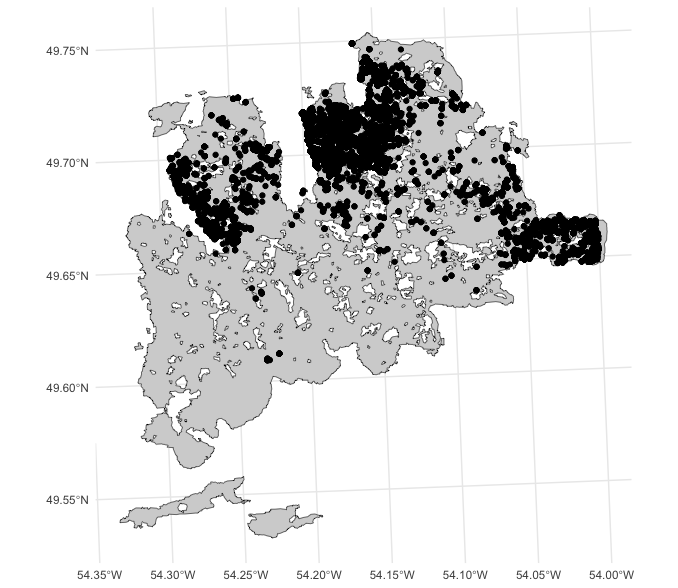

**Figure S2.** A plot of all fusion events for the winter period on Fogo Island from 2016 to 2025. Across all ten years (2016-2025), 11720 fusion events occurred in the winter season (Jan 1 to Mar 1) involving 43 individual caribou. A mean of 117.20 fusion events per winter, with approximately 54.51 events occurring per individual (as two individuals are involved in one fusion event) for each winter.

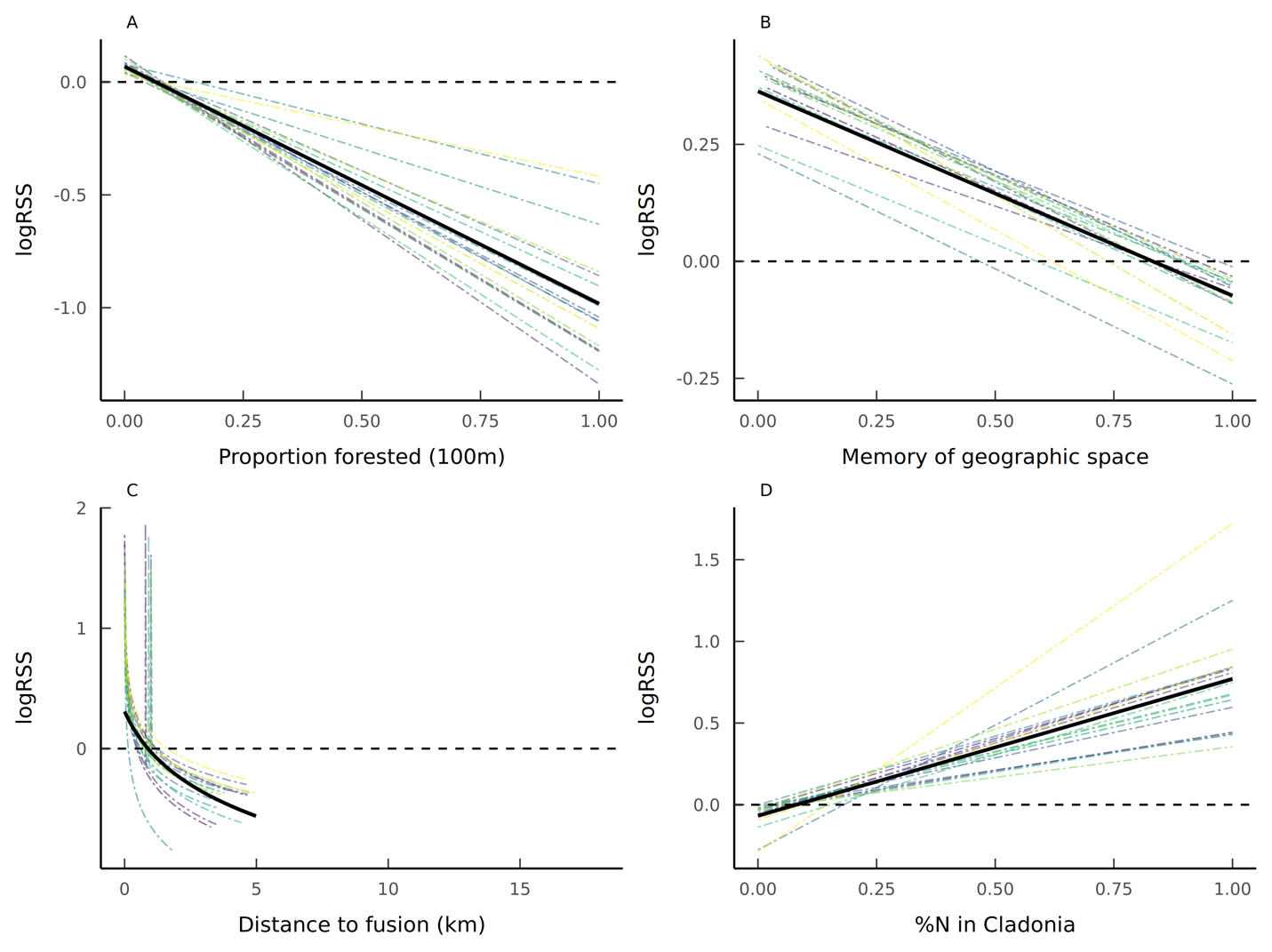

**Figure S3**. Relative selection strength (RSS) plots for proportion of forest (A) and long-term memory (B-D) with a utilization distribution restricted with a 50 % kernel. B shows the response to geographic memory alone, while C shows selection for previous fusion locations, and D shows selection for locations within long term memory that have higher *Cladonia* %N. The thick black line represents the population mean selection strength. Each individual is represented by their own dashed line, showing variation within the population. A value above 0 on the y-axis (logRSS) indicates selection for each parameter, when all other parameters are held at the mean. The dashed line at 0 on the y-axis also indicates at what value of the x-axis, selection changes to avoidance.

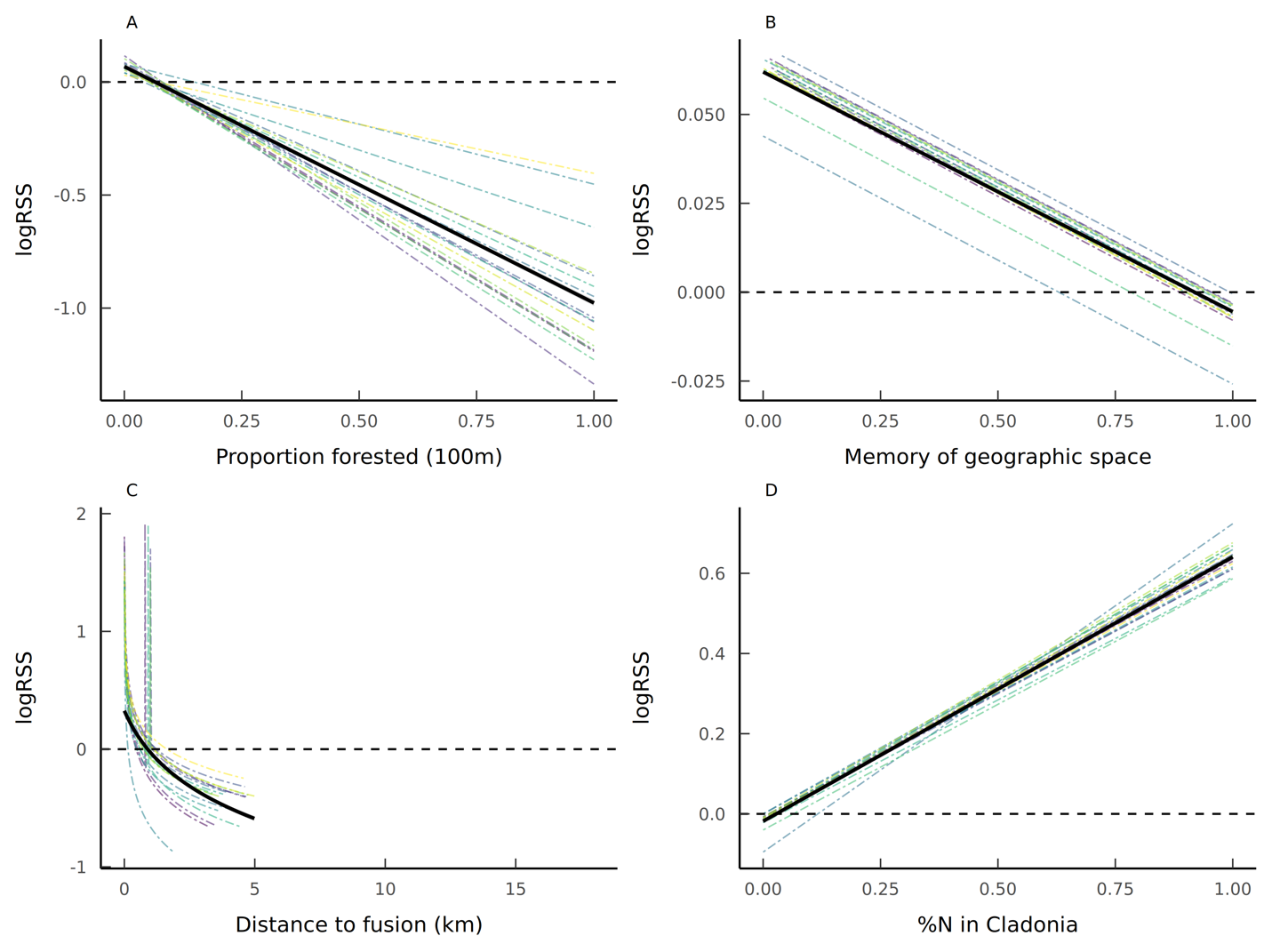

**Figure S4**. Relative selection strength (RSS) plots for proportion of forest (A) and long-term memory (B-D) with a utilization distribution restricted with a 25 % kernel. B shows the response to geographic memory alone, while C shows selection for previous fusion locations, and D shows selection for locations within long term memory that have higher *Cladonia* %N. The thick black line represents the population mean selection strength. Each individual is represented by their own dashed line, showing variation within the population. A value above 0 on the y-axis (logRSS) indicates selection for each parameter, when all other parameters are held at the mean. The dashed line at 0 on the y-axis also indicates at what value of the x-axis, selection changes to avoidance.

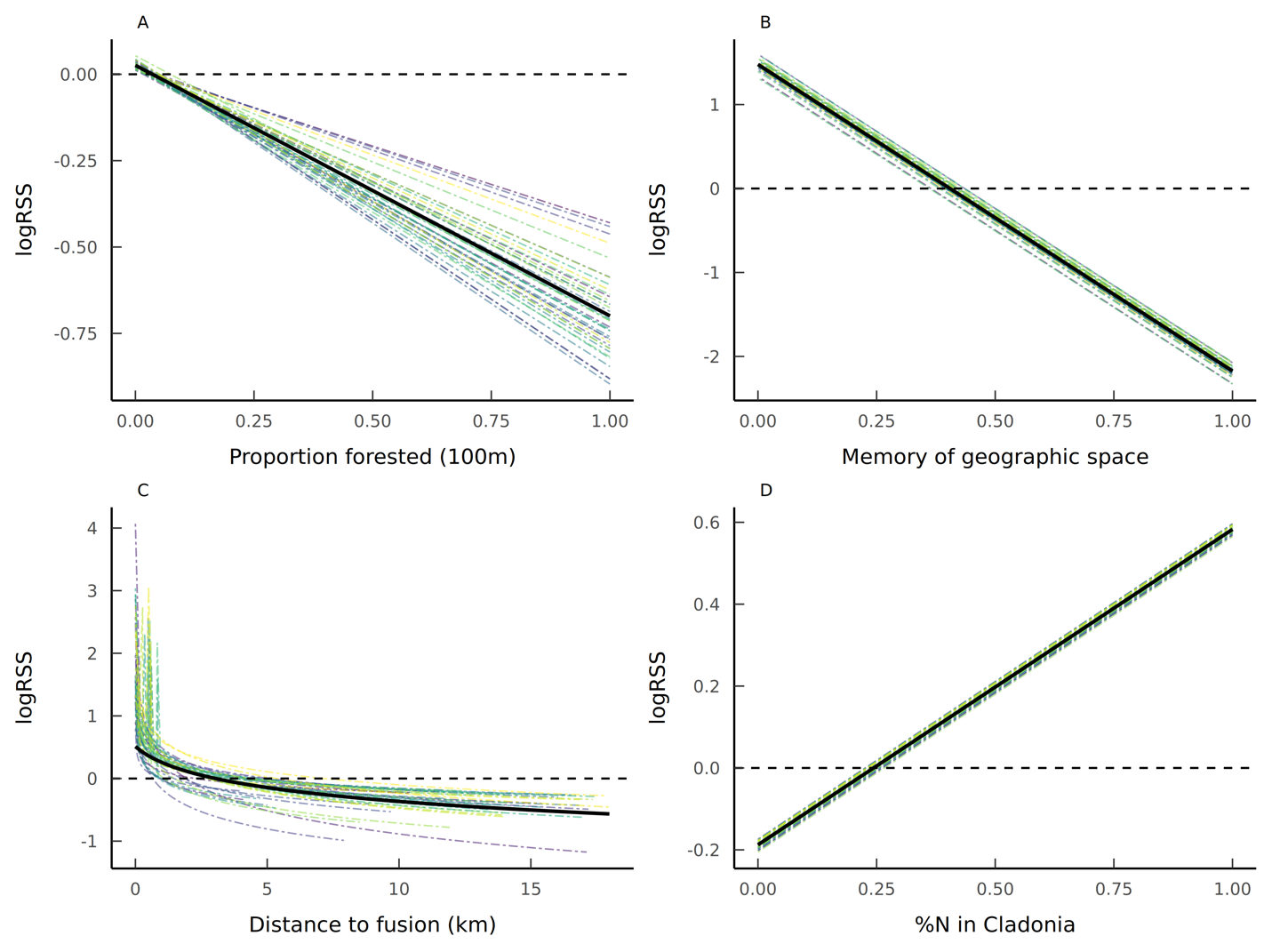

**Figure S5**. Relative selection strength (RSS) plots for proportion of forest (A) and short-term (3 day) memory (B-D). B shows the response to geographic memory alone, while C shows selection for previous fusion locations, and D shows selection for locations within short term memory that have higher *Cladonia* %N. The thick black line represents the population mean selection strength. Each individual is represented by their own dashed line, showing variation within the population. A value above 0 on the y-axis (logRSS) indicates selection for each parameter, when all other parameters are held at the mean. The dashed line at 0 on the y-axis also indicates at what value of the x-axis, selection changes to avoidance.

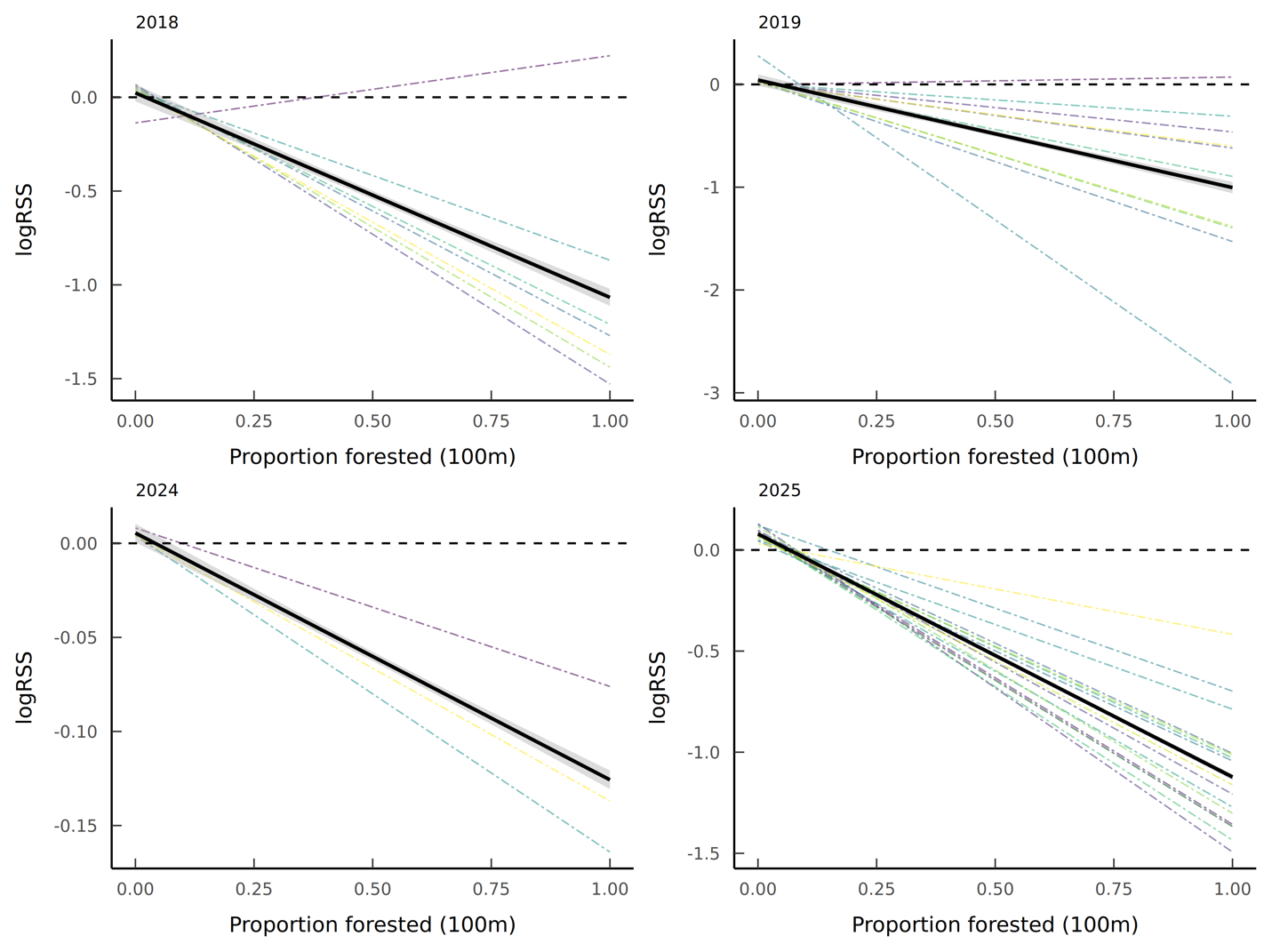

**Figure S6**. Relative selection strength (RSS) plots for proportion of forest in long-term memory models for years with the greatest sample size (2018, 2019, 2024, 2025). The thick black line represents the population mean selection strength. Each individual is represented by their own dashed line, showing variation within the population. A value above 0 on the y-axis (logRSS) indicates selection for each parameter, when all other parameters are held at the mean. The dashed line at 0 on the y-axis also indicates at what value of the x-axis, selection changes to avoidance.

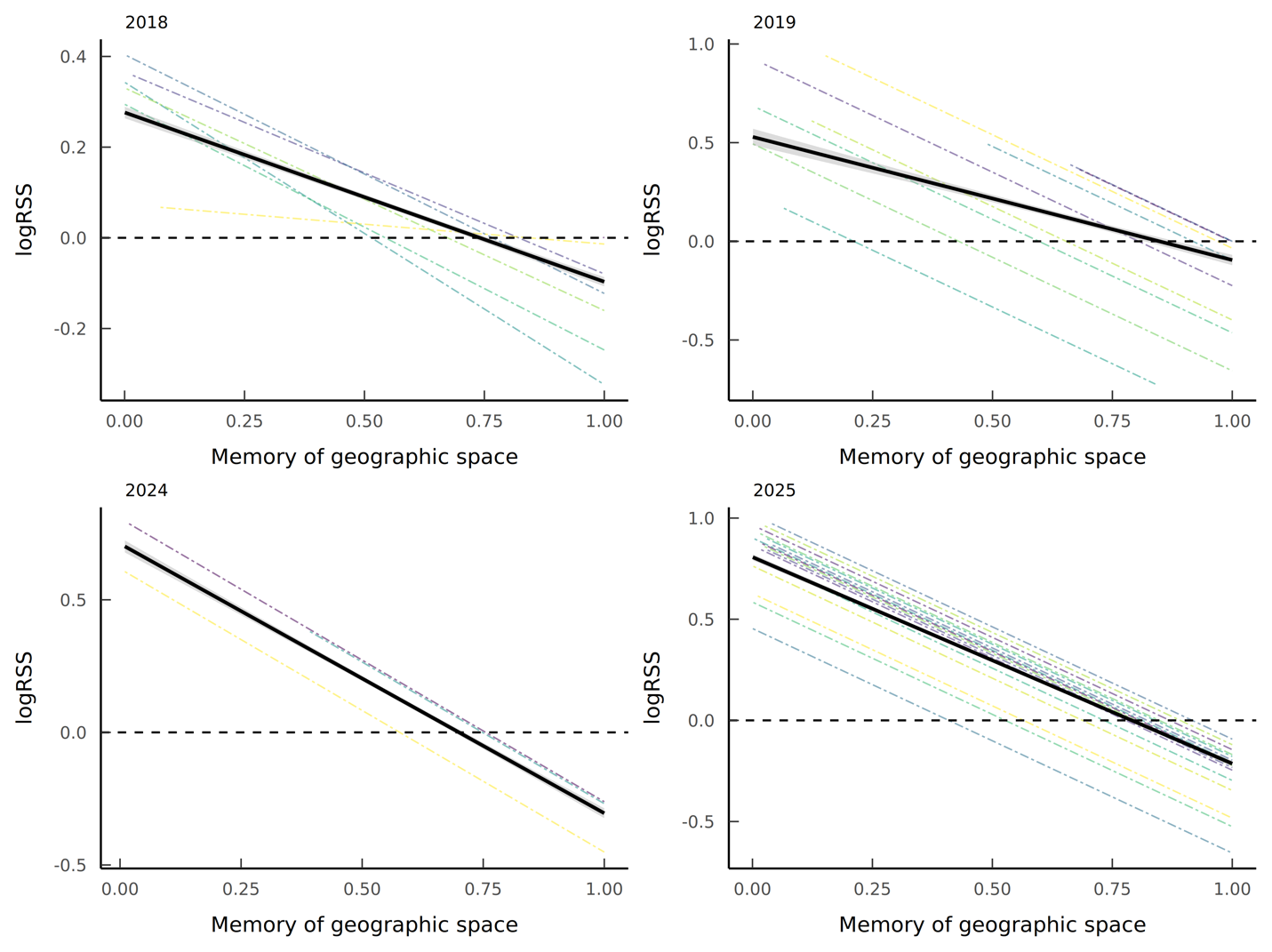

**Figure S7**. Relative selection strength (RSS) plots for geographic memory in long-term models for years with the greatest sample size (2018, 2019, 2024, 2025). The thick black line represents the population mean selection strength. Each individual is represented by their own dashed line, showing variation within the population. A value above 0 on the y-axis (logRSS) indicates selection for each parameter, when all other parameters are held at the mean. The dashed line at 0 on the y-axis also indicates at what value of the x-axis, selection changes to avoidance.

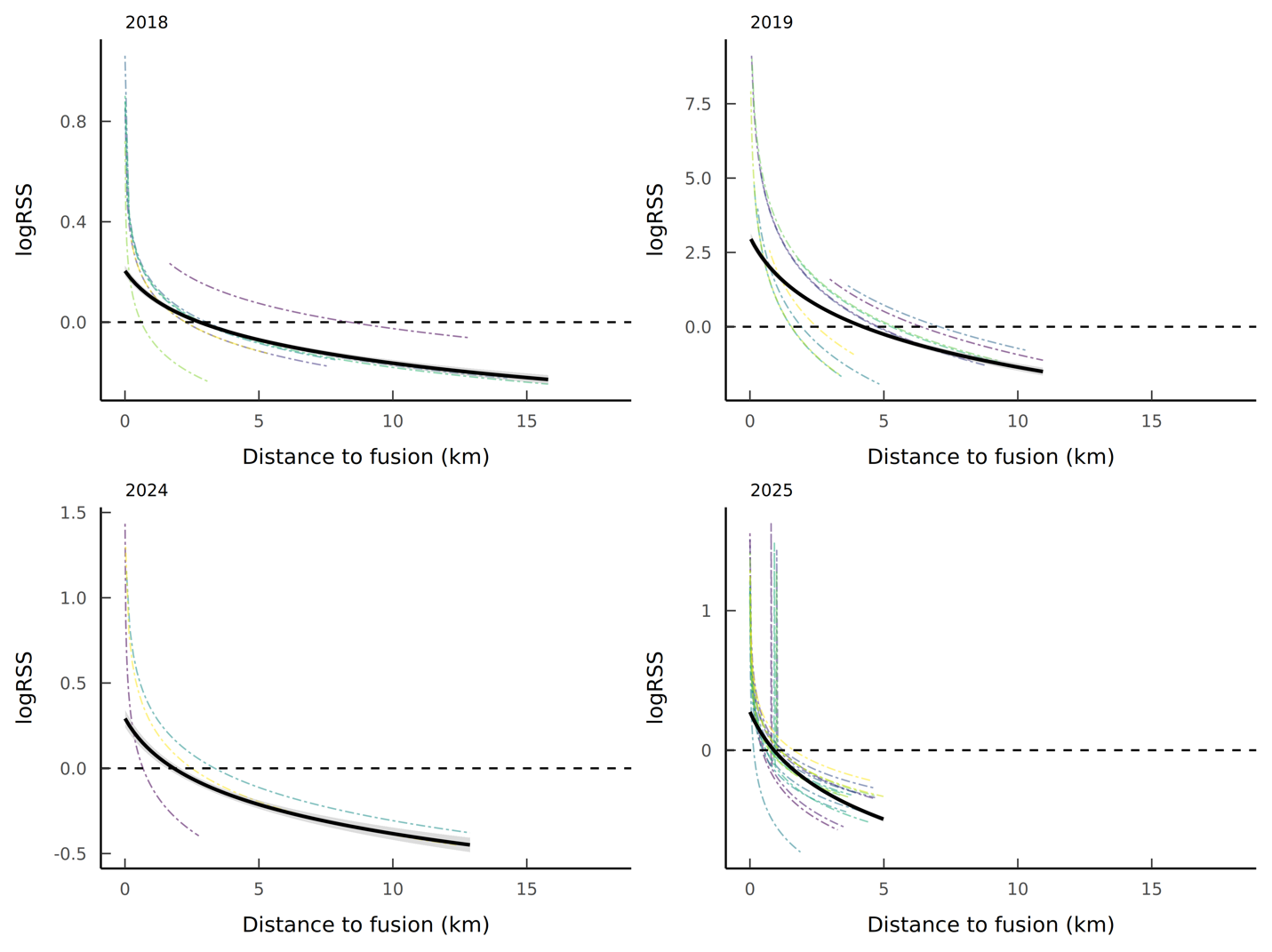
**Figure S8**. Relative selection strength (RSS) plots for social fusion in long-term memory models for years with the greatest sample size (2018, 2019, 2024, 2025). The thick black line represents the population mean selection strength. Each individual is represented by their own dashed line, showing variation within the population. A value above 0 on the y-axis (logRSS) indicates selection for each parameter, when all other parameters are held at the mean. The dashed line at 0 on the y-axis also indicates at what value of the x-axis, selection changes to avoidance.

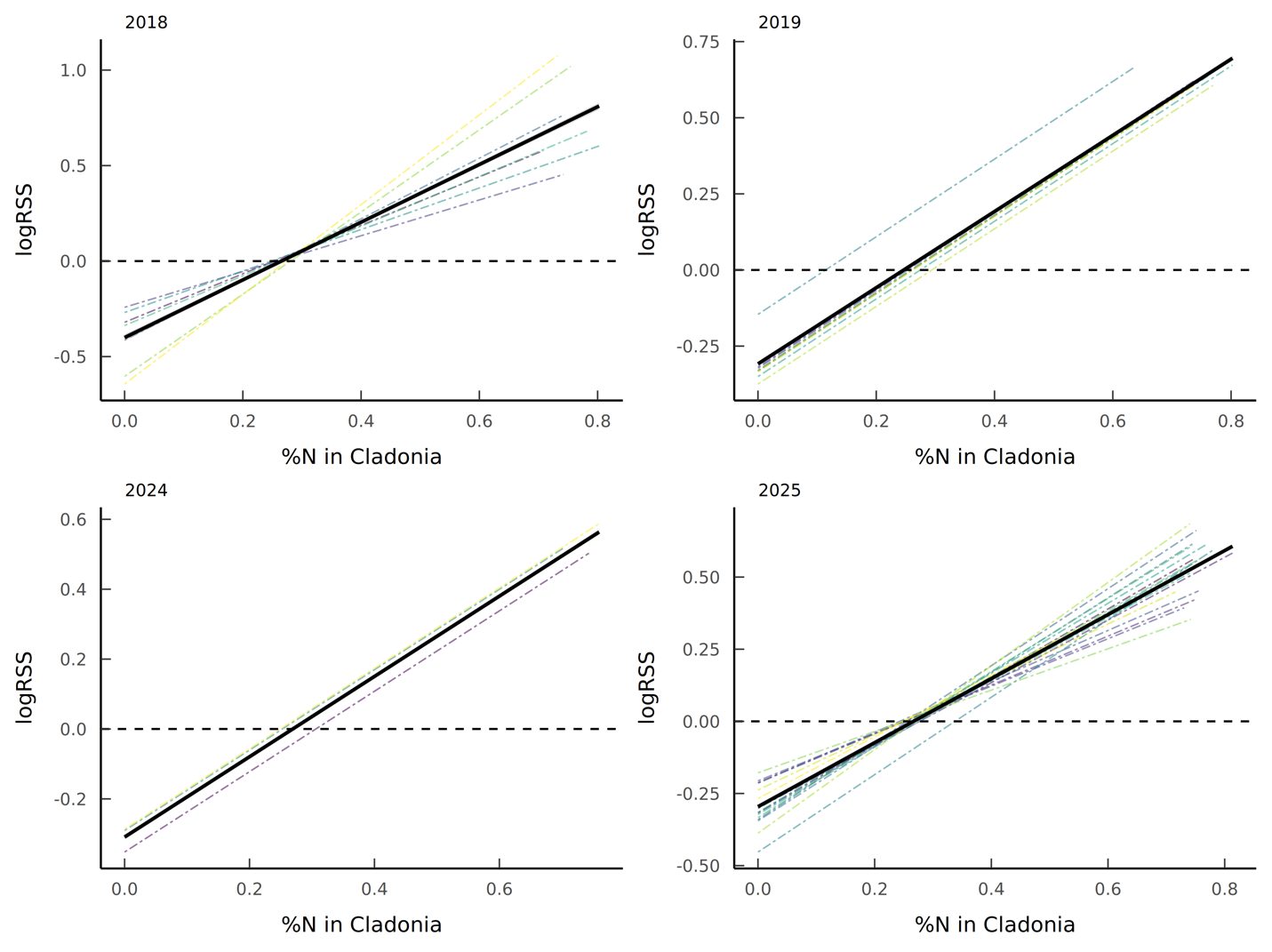

**Figure S9**. Relative selection strength (RSS) plots % N in *Cladonia* in long-term memory models for years with the greatest sample size (2018, 2019, 2024, 2025). The thick black line represents the population mean selection strength. Each individual is represented by their own dashed line, showing variation within the population. A value above 0 on the y-axis (logRSS) indicates selection for each parameter, when all other parameters are held at the mean. The dashed line at 0 on the y-axis also indicates at what value of the x-axis, selection changes to avoidance.

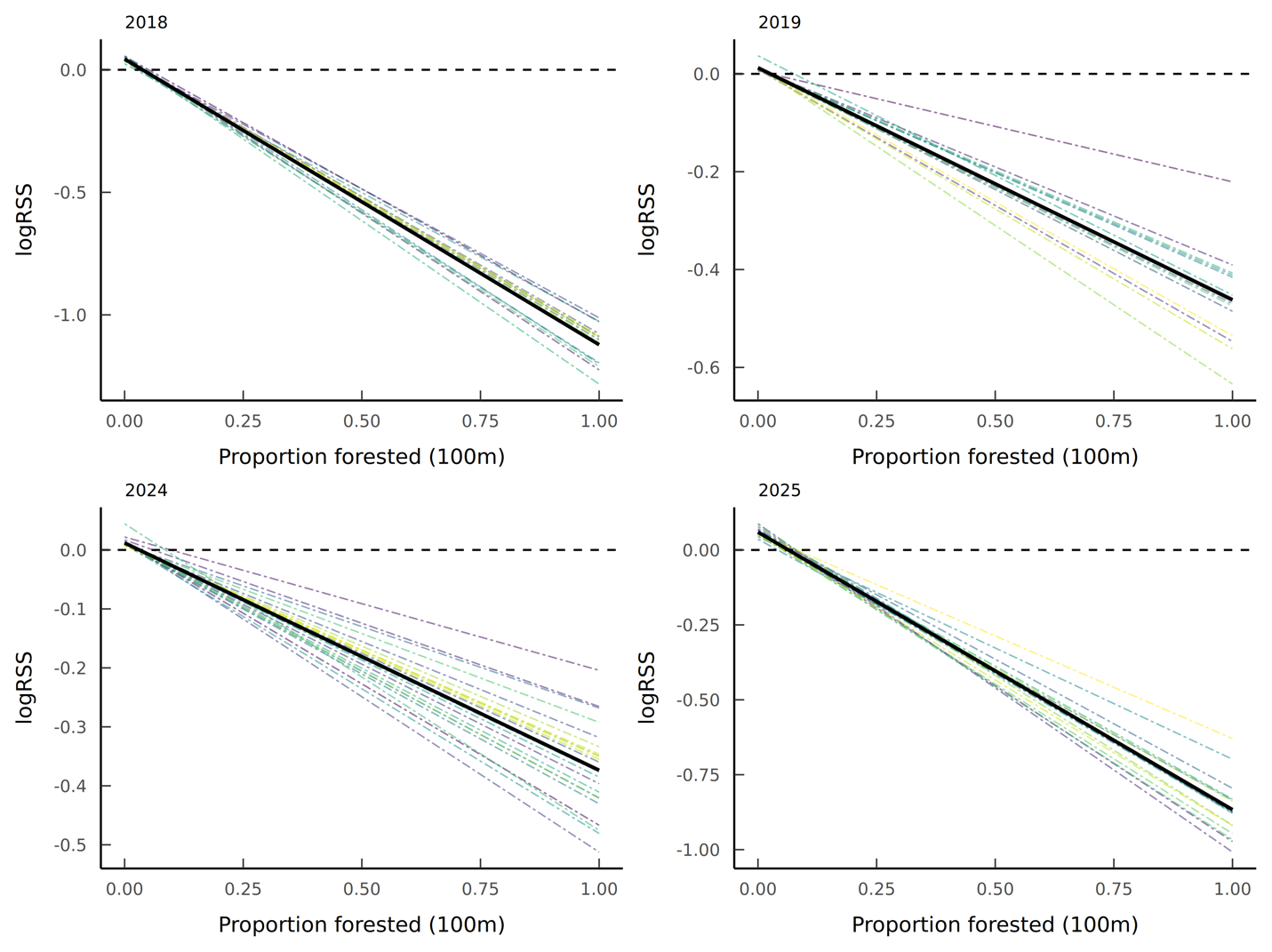

**Figure S10**. Relative selection strength (RSS) plots for proportion of forest in short-term memory models for years with the greatest sample size (2018, 2019, 2024, 2025). The thick black line represents the population mean selection strength. Each individual is represented by their own dashed line, showing variation within the population. A value above 0 on the y-axis (logRSS) indicates selection for each parameter, when all other parameters are held at the mean. The dashed line at 0 on the y-axis also indicates at what value of the x-axis, selection changes to avoidance.

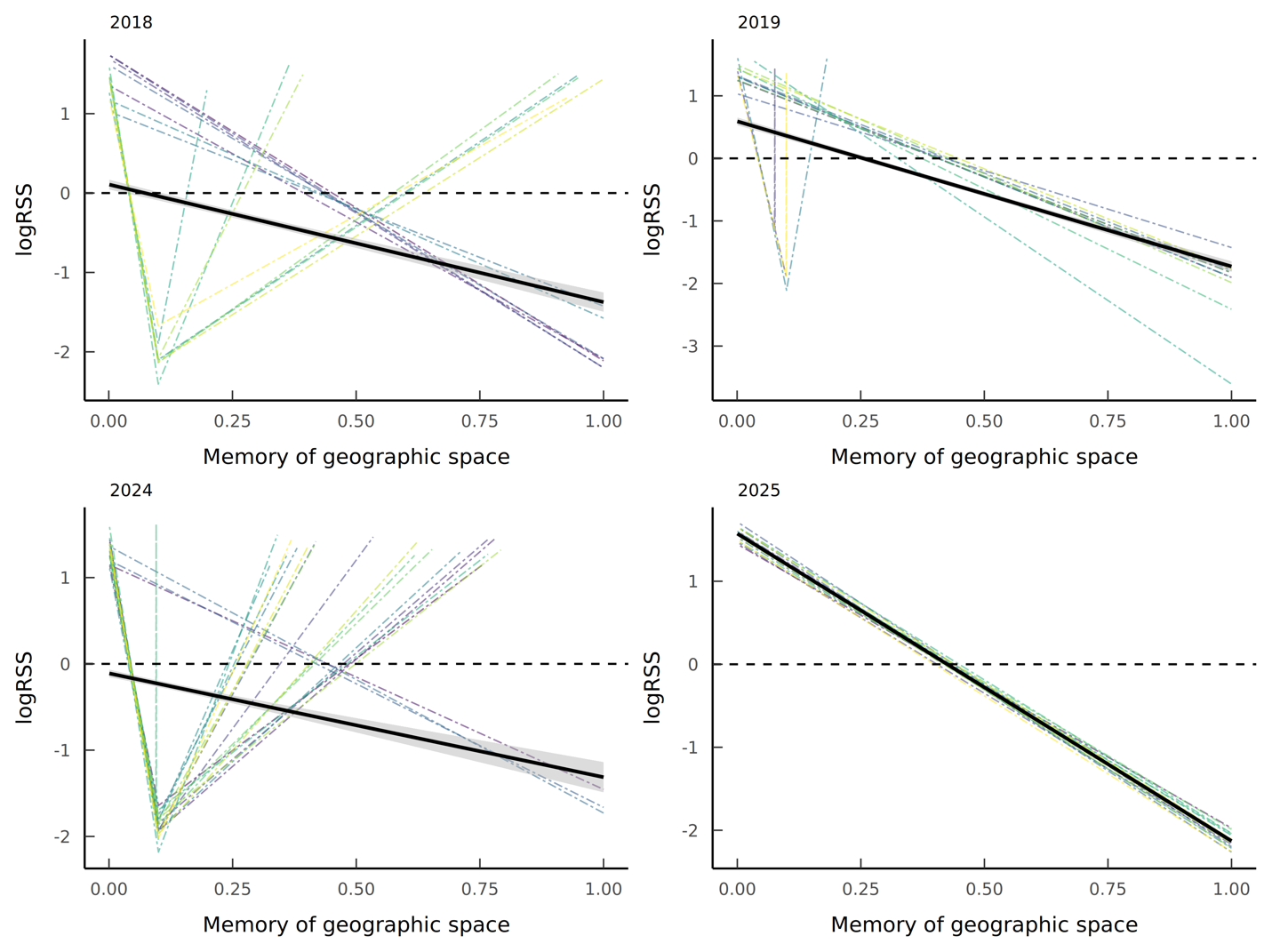
**Figure S11**. Relative selection strength (RSS) plots for geographic memory in short-term memory models for years with the greatest sample size (2018, 2019, 2024, 2025). The thick black line represents the population mean selection strength. Each individual is represented by their own dashed line, showing variation within the population. A value above 0 on the y-axis (logRSS) indicates selection for each parameter, when all other parameters are held at the mean. The dashed line at 0 on the y-axis also indicates at what value of the x-axis, selection changes to avoidance.

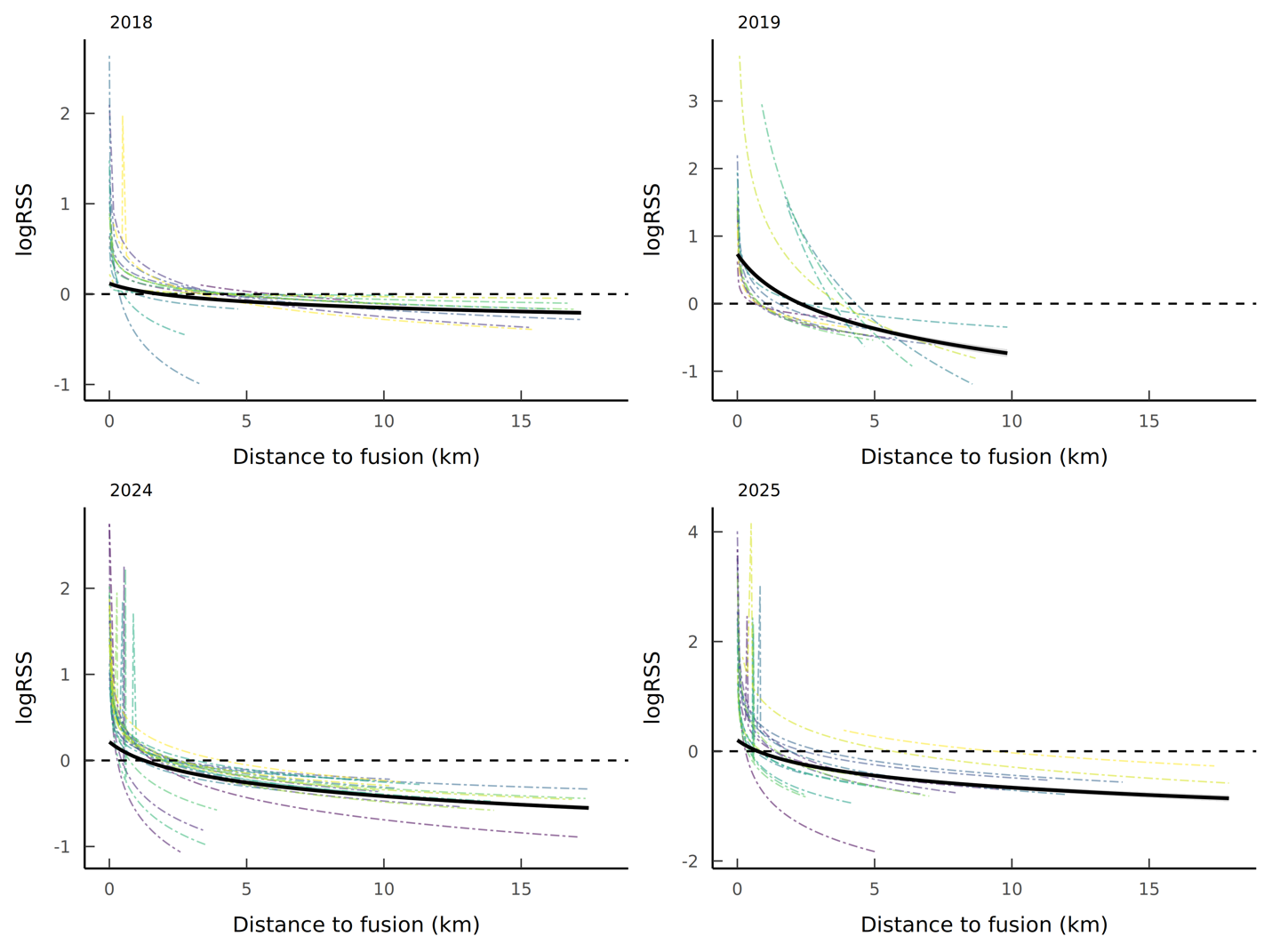

**Figure S12**. Relative selection strength (RSS) plots for social fusion in short-term memory models for years with the greatest sample size (2018, 2019, 2024, 2025). The thick black line represents the population mean selection strength. Each individual is represented by their own dashed line, showing variation within the population. A value above 0 on the y-axis (logRSS) indicates selection for each parameter, when all other parameters are held at the mean. The dashed line at 0 on the y-axis also indicates at what value of the x-axis, selection changes to avoidance.

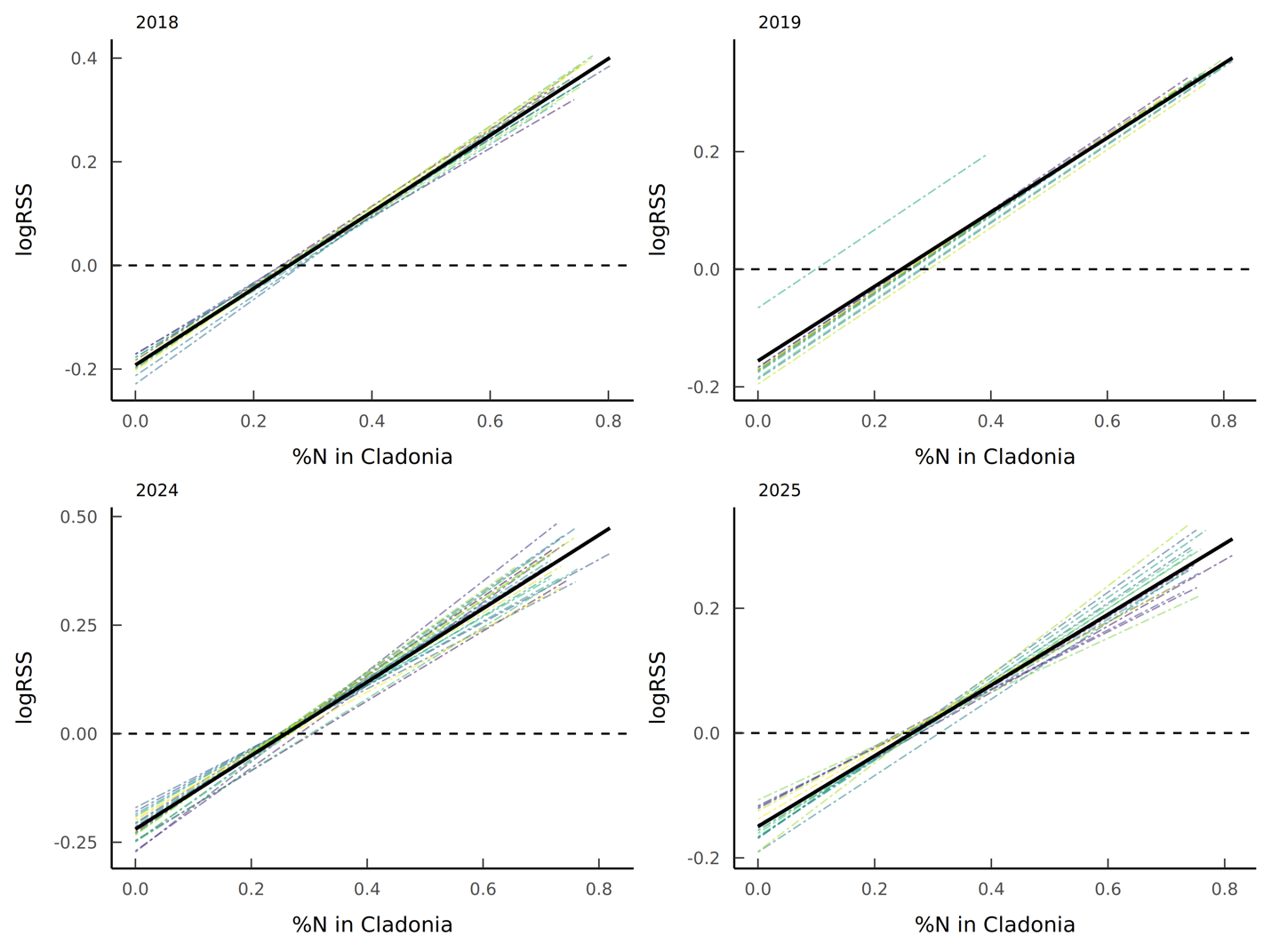

**Figure S13**. Relative selection strength (RSS) plots for %N of *Cladonia* in short-term memory models for years with the greatest sample size (2018, 2019, 2024, 2025). The thick black line represents the population mean selection strength. Each individual is represented by their own dashed line, showing variation within the population. A value above 0 on the y-axis (logRSS) indicates selection for each parameter, when all other parameters are held at the mean. The dashed line at 0 on the y-axis also indicates at what value of the x-axis, selection changes to avoidance.
